# A system for high-throughput axonal imaging of induced pluripotent stem cell-derived human i^3^Neurons

**DOI:** 10.1101/2025.11.22.689945

**Authors:** Stella A.C. Whittaker, Elizabeth D. McKenna, Stephanie L. Sarbanes, Michael S. Fernandopulle, Michael E. Ward, Antonina Roll-Mecak

**Author notes:** Corresponding author: Antonina Roll-Mecak, Cell Biology and Biophysics Unit, Porter Neuroscience Research Center, National Institutes of Health, Building 35, Room 3B-203, 35 Convent Drive, MSC 3700, Bethesda, MD 20892-3700.

## Abstract

Neurons have long, thin axons and branched dendritic processes which rely on an extensive microtubule network that functions as a cellular scaffold and substrate for cargo transport. Microtubule defects are a defining pathological feature of neurological disorders. The highly arborized, long, polarized neuronal processes pose challenges for imaging-based assays. Available methods use either dispersed cultures, which are inefficient for compartment-specific analyses, or microfluidic chambers, which allow clear separation of somatodendritic and axonal compartments but are expensive and difficult to maintain. Here, we introduce an “i^3^Neurosphere” culture model of induced pluripotent stem cell (iPSC)-derived human cortical i^3^Neurons that enables high-throughput imaging of hundreds of axons without specialized equipment. We characterize neurite outgrowth, polarization, microtubule dynamics, and motility of diverse cargo, providing a reference for future work on microtubule processes in this system. The high-throughput compartment-specific imaging we present, combined with facile genetic engineering in i^3^Neurons provides a powerful tool to study human neurons.

**SIGNIFICANCE STATEMENT:** Human neurons are difficult to study due to limited access to tissue and technical challenges in existing *in vitro* models of axonal transport.

We developed *i^3^Neurospheres*, a simple and scalable 3D culture system of human iPSC-derived neurons that enables high-throughput imaging of axonal outgrowth, microtubule dynamics, and intracellular transport.

This platform provides an accessible, reproducible method for investigating neuronal function and disease mechanisms, offering broad utility for neuroscience research and preclinical drug screening.

## INTRODUCTION

Nervous system function relies on the precise spatiotemporal targeting of neuronal processes and the efficient trafficking of cargo to support these processes, enabling their constant remodeling in response to environmental cues. Key to nervous system wiring is the microtubule array, a cytoskeletal network of interconnected filaments that mechanically supports process extension and serves as tracks for intracellular cargo transport by molecular motors. Defects in microtubules and microtubule-dependent trafficking are a salient hallmark of neurological disease (Ioannou *et al*., 2019; Sleigh *et al*., 2019), which confers an enormous toll in human suffering and economic burden. Great strides in understanding fundamental neuronal biology are required to make meaningful progress towards treating them. Microtubules are composed of α/β-tubulin heterodimers stacked longitudinally into polar protofilaments that laterally associate to form rigid yet dynamic, hollow cylindrical polymers. Microtubules in the axon are almost uniformly plus-end out, while microtubules in dendrites exhibit mixed polarity. The distinct tubulin genes that compose microtubules combined with the myriad post-translational modifications that chemically modify tubulin are together referred to as the “tubulin code” and comprise the most fundamental layer of microtubule regulation. A second layer of regulation is achieved with microtubule associated proteins (MAPs) (McKenna *et al*., 2023). Beyond this, motors and motor adaptors that link motors to diverse cargo are the effectors that drive cellular transport. The kinesin superfamily of molecular motors mediates plus-end directed, and thus anterograde axonal transport, away from the soma and toward the distal tips of neurites to supply essential cargo to synapses. The molecular motor dynein mediates minus-end directed, and thus retrograde, axonal transport, away from neurite tips and toward the soma, carrying back trophic support and signaling (Guedes-Dias and Holzbaur, 2019). Together, tubulin modifications, MAPs, motors and motor adapters confer interrelated feedback to coordinate the building and maintenance of functionally and morphologically distinct axonal and dendritic compartments. Mutations in all these components, the αβ-tubulin building blocks, tubulin modification enzymes, MAPs, molecular motors, and motor adapters cause neuronal dysfunction in diverse model systems as well as in humans (McKenna *et al*., 2023).

Studying the human neuronal microtubule cytoskeleton is challenging because the human central nervous system (CNS) is inaccessible and because neurons are post-mitotic, and thus not a renewable tissue source. Therefore, the field has largely relied on *in vitro* models using cortical or hippocampal neurons dissected from mouse brain (Boecker *et al*., 2020). However, recent years have seen an ascendence in using human stem-cell derived neurons. i^3^Neurons (inducible, integrated, isogenic) are derived from a well-characterized iPSC line and encode a doxycycline-inducible neurogenin (NGN2) expression cassette integrated at the AAVS1 safe-harbor locus that allows for highly reproducible, stereotyped neuron generation at scale (Fernandopulle *et al*., 2018), which is advantageous compared to the expensive, relatively finite, and technically challenging brain dissections necessary for primary neuron preparations or the more variable small molecule-based differentiation iPSC protocols (Qi *et al*., 2017; Fernandopulle *et al*., 2018). Additionally, gene editing is technically simpler in iPSCs and enables easy tagging with fluorescent proteins for live-cell imaging, e.g. the imaging of neuronal cargo or molecular motors, at their endogenous expression levels (Roberts *et al*., 2017). Human iPSCs, as opposed to those that are of non-human origins, are also particularly advantageous due to the vast genomic non-coding regions that are not conserved between species, and whose contributions to physiological function or human disease are unknown (Ichida and Kiskinis, 2015). We note that using i^3^Neurons results in the loss of the 3-dimensional physiological and cellular context in which human neurons develop *in vivo*, however, *in vivo* animal models lack the advantages of the human genetic context previously listed, are more expensive, and can lack spatiotemporal imaging resolution.

Beyond the potential for fundamental neurobiological discovery, the human iPSC-derived neuronal model system can be applied to disease research with several outstanding advantages. (*i*) The wild-type neurons can be compared to mutant neurons at endogenous expression levels of mutant proteins through CRISPR/Cas9-mediated generation of knock-in lines. (*ii*) The dox-inducible NGN2 gene can easily be introduced into patient iPSCs allowing for the study of disease in patient-specific samples as well as the study of sporadic disease, which constitutes a vast subset of neurological diseases and can only be modeled using patient tissue (Ichida and Kiskinis, 2015). (*iii*) Most significantly, failed drug trials based on animal models suggest that an important therapeutic approach will be to test drugs in patient models prior to expensive clinical trials (Cudkowicz *et al*., 2006; Blesa and Przedborski, 2014; Kang *et al*., 2021; Azkona and Sanchez-Pernaute, 2022).

Previous investigations related to microtubule-based transport in i^3^Neurons have shown them to be versatile and powerful for cargo transport assays (Liao *et al*., 2019; Boecker *et al*., 2020).These investigations relied on either dispersed low-density i^3^Neuron cultures (Boecker *et al*., 2020) or microfluidic devices that guide neurite growth through microfabricated channels to allow separation between the axonal and somatodendritic compartments (Taylor *et al*., 2003; Park *et al*., 2006; Poon *et al*., 2013; Neto *et al*., 2016). Microfluidic devices are cost-prohibitive for many research endeavors and can require extensive optimizations. Dispersed cultures largely restrict image acquisition to one axon per image, severely limiting data collection. Moreover, they rely on morphological characterization for axon identification, which is cumbersome. In contrast, the well-established neuronal explant model, often derived from rodent dorsal root ganglia, has enabled foundational research on axon guidance, cargo transport, regeneration and response to guidance cues in a controlled environment that allows rapid analysis of hundreds of axons in a physiologically relevant context (Levi-Montalcini, 1987; Baas and Black, 1990; Metin *et al*., 1997; Bilsland *et al*., 1999). However, as dissected primary tissue, explants are subject to the same limitations as other primary tissue as discussed above. Here, we combine the advantages of explant methodology with human iPSC-derived neurons to generate “i^3^Neurospheres” that enable facile visual separation of neuronal compartments, and cost-effective, high-throughput imaging of thousands of i^3^Neuron axons simultaneously (Dargan *et al*., 2024; De Pace *et al*., 2024). Using these i^3^Neurospheres, we characterize axonal outgrowth, microtubule dynamics, and motor transport of various cargoes needed for neuronal survival and signaling, and benchmark them against those measured in dispersed cultures. Our work introduces a powerful tool to study axonal transport in human i^3^Neurons that can be easily applied in any lab without the need for specialized reagents or equipment. Our quantitative analysis of microtubule dynamics and transport kinetics of diverse cellular cargo provides a baseline reference for future studies on cytoskeletal function in i^3^Neurons and mechanisms of neurological disease.

## RESULTS

### i^3^Neurosphere Projections Extend Radially in a Highly Stereotyped Manner

Axonal outgrowth from explants is an invaluable and established tool to study axonal extension, cargo transport, regeneration and response to guidance cues in a controlled environment that allows easy simultaneous imaging of hundreds of axons (Levi-Montalcini, 1987; Baas and Black, 1990; Metin *et al*., 1997; Bilsland *et al*., 1999). To combine the strength of such a system with the advantages of the i^3^Neurons, we devised a protocol for generating neurospheres by combining the previously described dispersed culture i^3^Neuron protocol (Fernandopulle *et al*., 2018) for differentiation of dox-inducible NGN2 neurons (Wang *et al*., 2017) (Methods) with an established method for generating cardiac organoids (Hoang *et al*., 2018). We plated ∼10,000 iPSCs stably expressing mScarlet for cellular visualization (Methods) into a low-attachment 96 well plate with doxycycline-containing neuronal induction media. Plates were spun and wells were checked to verify that the iPSCs had assembled into small spheroids. After 4 days of neuronal induction, these neural spheroids, termed i^3^Neurospheres, were manually picked using a P200 pipette with wide-orifice pipette tips and replated onto a glass-bottom dish coated with poly-L-ornithine and laminin (Methods, Figure 1A). In contrast to the i^3^Neurons in dispersed culture, which display a heterogenous, random outgrowth of neuronal processes (Figure 1B), i^3^Neurosphere outgrowth is highly reproducible, with neuronal projections growing radially from the center (Figure 1C). We used the i^3^Neurosphere radius as a metric for the average neurite outgrowth of thousands of neurites (Figure 1E). Their average radius changed from 0.70 ± 0.03 mm at day 5 (D5) to 1.37 ± 0.02 mm at D7 and 2.47 ± 0.97 mm at D11 of differentiation (Figure 1D). This time course reflects the period of rapid neurite outgrowth prior to contact with the walls of the culture dish. We note that we uniformly observed a swirling radial morphology in i3Neurospheres that intensifies over time, which is likely attributable to media changes, despite extreme caution taken to change media very gently. Overall, these data demonstrate a cost-effective, reproducible method of assessing outgrowth of a large population of neurites from human iPSC-derived i^3^Neurons.

**Figure 1.**
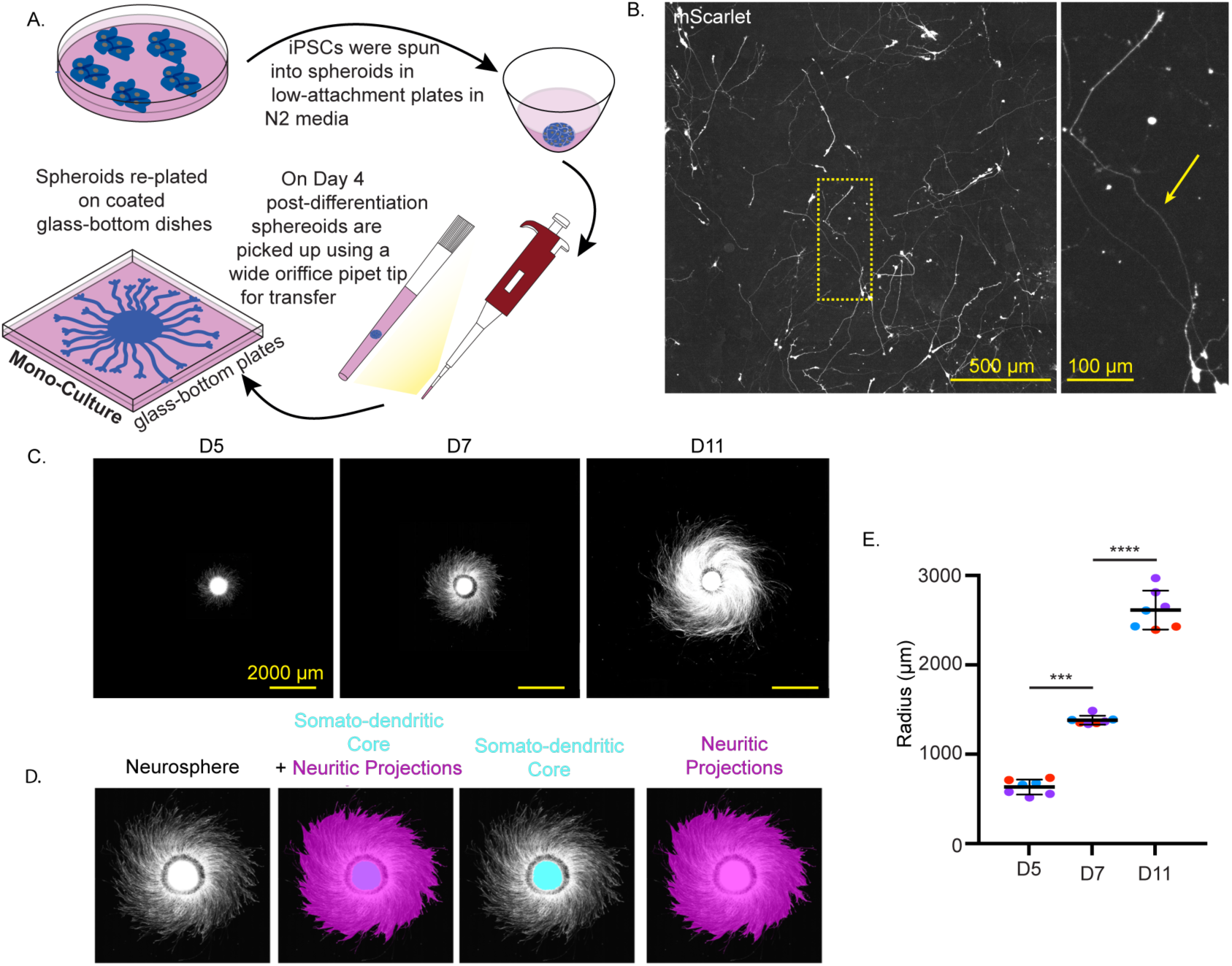
Neurospheres of iPSC-derived i^3^Neurons are a versatile tool for monitoring neuronal outgrowth. (A) Schematic showing method for generating i^3^Neurospheres. (B) Image of dispersed culture of mScarlet-expressing i^3^Neurons at D11. Arrow indicates the axon. (C) Images of neuritic outgrowth in mScarlet-expressing neurospheres as a function of differentiation time. (D) Image of neuritic outgrowth, highlighting the inner somatodendritic core (cyan) and the boundary of neuritic projections (magenta) after maximum intensity projection and thresholding. (E) Radius of individual neurospheres as a function of differentiation stage. Each circle represents an individual neurosphere; colors represent independent differentiations. Bars represent mean and S.D., ***, p>0.001, ****, p>0.0001 by unpaired t-test. N = 7 neurospheres from three independent differentiations.

### Distal i^3^Neurosphere Projections are Axons

The microtubule array reflects the functional specificity of the neuron, including compartment specification for the dendritic arbor and the axon. MAP2 and tau are two canonical neuron-specific MAPs. MAP2 is restricted to the somatodendritic compartment, while tau is distributed throughout the neuron (Monroy *et al*., 2020). To distinguish the identity of i^3^Neurosphere projections, we used immunocytochemistry with antibodies specific to MAP2 and tau. Image analysis over a time course of early differentiation shows a somatodendritic sphere radius of ∼250μm and dendritic processes that terminate ∼200 μm (D5), ∼430 μm (D7) and ultimately plateau at ∼850 μm (D9 and 11) from the center of the somatodendritic spheroid core (Figures 2A, B). These data indicate that imaging regions of neurites distal to these distances at each time point ensures selective analysis of axonal compartments in the i^3^Neurospheres. This can easily be achieved by imaging regions less than ∼500μm from the growing neurite edge before D11, and ∼1 mm after D11 (Figure 2B). Axonal identity in these regions was further validated using an antibody for axon-specific neurofilaments (Ulfig *et al*., 1998), which displayed robust labeling of neuritic processes with the vast majority already extending beyond the boundary of the MAP2 signal by D5 (Fig. S1). In conclusion, our immunofluorescence analysis shows that i^3^Neurosphere assays allow the easy identification of hundreds of axons for efficient imaging.

**Figure 2.**
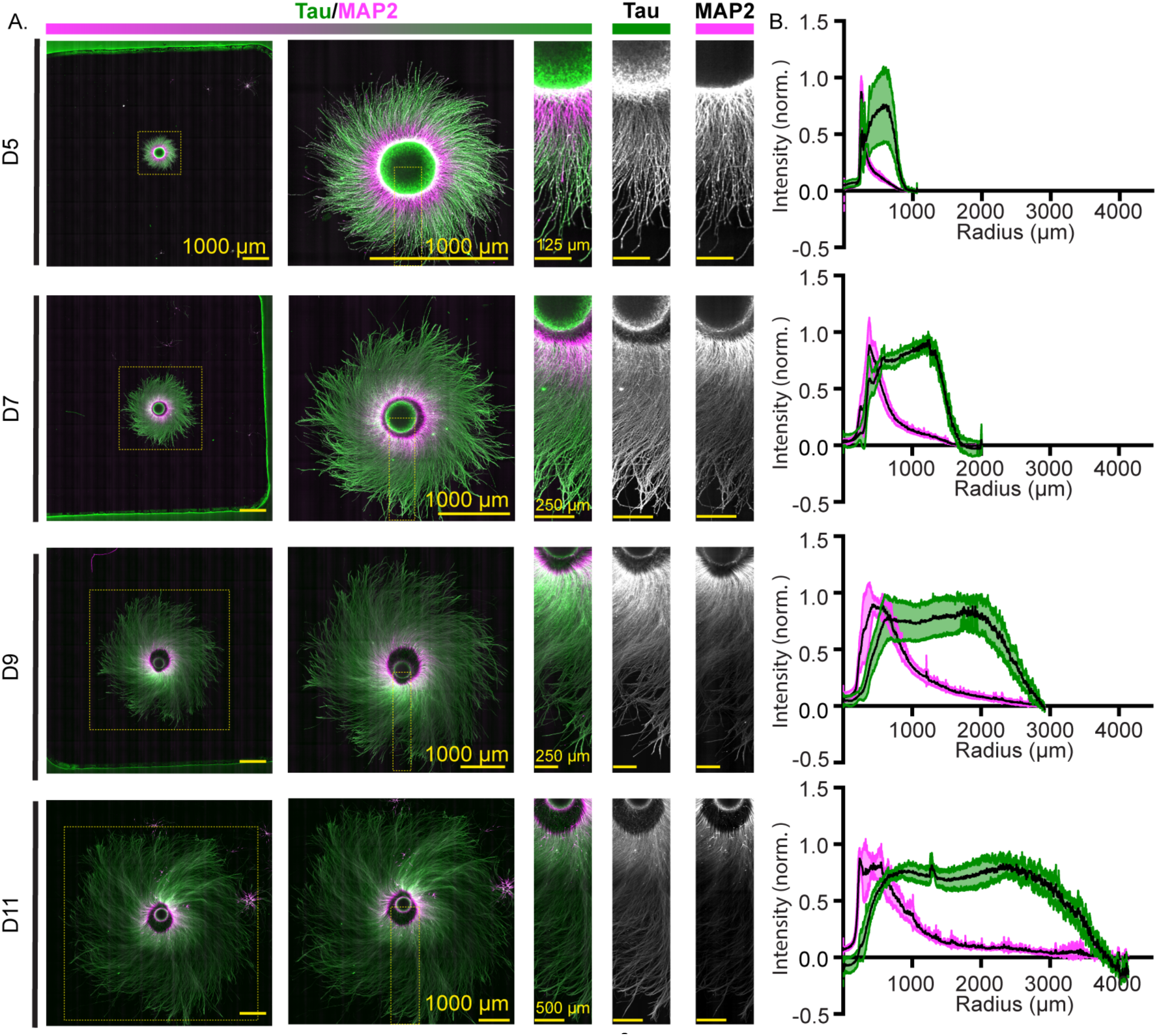
Neuronal polarity markers in i^3^Neurospheres define dendritic and axonal compartment boundaries. (A) Representative images of neurospheres immunolabeled for MAP2 (magenta) and tau (green). (B) Line plots along the neurosphere radius (from center to periphery) showing average intensity of MAP2 (magenta) and tau (green), each normalized to their respective highest intensity, for D5, D7, D9 and D11. Black line, average, green and magenta shaded areas, S.D. for MAP2 and tau intensity, respectively; N = 8 neurospheres from two independent differentiations.

Neurons are primarily signaling cells in which axons transmit, and dendrites receive signals. Microtubules are overwhelmingly oriented plus-end out in the axon, and adopt mixed orientation in dendrites, reflecting the functional distinctiveness of these two compartments (Baas *et al*., 1988). Having established axonal identity, we next sought to characterize microtubule growth trajectory and dynamics in i^3^Neurosphere axons. Using mScarlet-expressing iPSCs that also stably express GFP-tagged End Binding Protein 1 (EB1), we tracked growing microtubules. EB1 recognizes the ends of growing microtubules, giving it the appearance of a “comet” (Honnappa *et al*., 2009) (Methods; Figure 3A; Supplemental Movie 1). We analyzed EB1 trajectories in the distal ∼ 200 μm of neurosphere axons and found that at D5 71% and 29% of EB comets were anterograde and retrograde, respectively. Notably, the proportion of anterograde comets increases monotonically with differentiation reaching 75% at D7, and 86% at D11 (Figure 3C), with ∼80% of neurite having only anterograde comets at D11, consistent with previous reports that microtubule orientation becomes increasingly plus-end-out during axonal development in primary neurons (del Castillo *et al*., 2015; Jakobs *et al*., 2022). These results are in line with previous reports in dispersed i^3^Neuron cultures that showed that at a later timepoint (D21), 99% of EB1 comets in axons grow in the anterograde direction (Boecker *et al*., 2020). We also quantified axonal microtubule growth speed using EB1 and determined an average instantaneous velocity that stayed constant throughout differentiation, from 0.11 ± 0.05 μm/s at D5 to 0.11 ± 0.07 μm/s for D7, and 0.1 ± 0.07 μm/s for D11 (Figure 3D). These microtubule growth speeds are in line with previous reported values of 0.15 μm/s in dispersed i^3^Neuron cultures obtained by tracking MACF43, another protein that tracks the ends of growing microtubules (Radwitz *et al*., 2022). The relatively uniform growth speeds we record are also consistent with the linear increase in neurite outgrowth we observe across our time course (Figure 1E).

**Figure 3.**
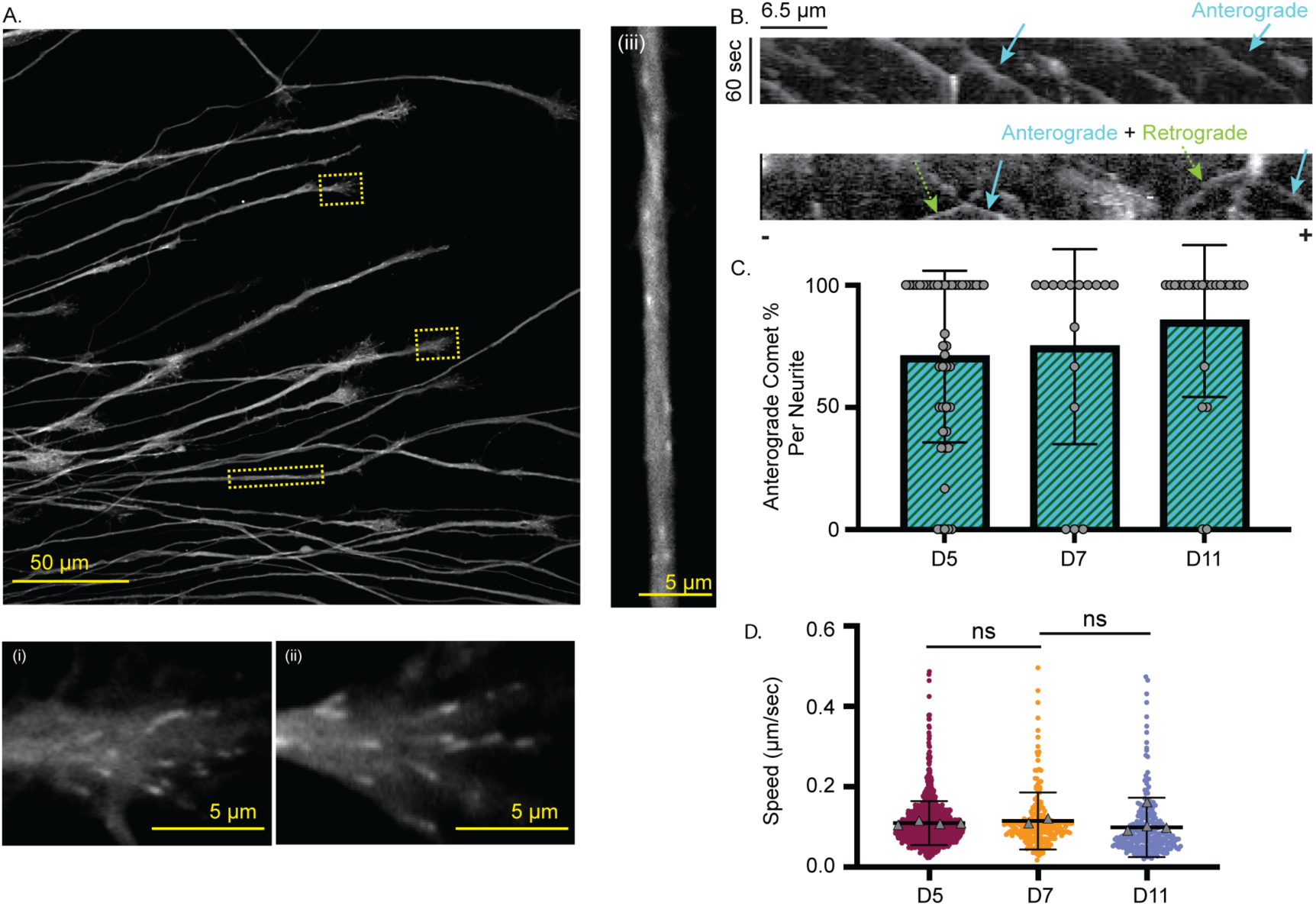
Microtubule polarity and orientation over time in i^3^Neurosphere axons. (A) Images of EB1-GFP in distal i^3^Neurosphere axons. Dashed lines mark insets showing two growth cones at higher magnification in panels (i) and (ii) and a segment of a distal neurite in panel (iii). (B) Kymograph of EB1-GFP microtubule comets in distal axons. Top panel, kymograph showing exclusively anterograde comets. Bottom panel, kymograph showing one or more retrograde comets. Arrowheads highlight anterograde (blue) and retrograde (green) comets with – and + denoting direction of neurite growth from soma to growth cone, respectively. (C) Percentage of anterograde (green hatched) comets per neurite as a function of differentiation time. Bars represent mean and S.D., each circle represents an individual neurite, D5: N = 46, D7: N = 15, D11: N = 23 neurites from two independent differentiations. (D) EB1 comet speeds in i^3^Neurosphere growth cones as a function of differentiation time. Each circle represents a comet, triangles represent mean speed from each individual neurosphere, bars represent mean and S.D., ns, not significant; ****, p < 0.0001, by Kruskal-Wallis test. D5, N = 1136, D7, N = 210, and D11, N = 253 comets from 60, 21, 35 growth cones respectively from two biological replicates from two independent differentiations.

### Longitudinal analysis of mitochondria motility in i^3^Neurosphere axons

Microtubule-dependent organelle transport is essential to neuronal function. Mitochondria are transported throughout axons and dendrites and dock at pre-synapses and synapses to provide energy and to buffer calcium that supports connectivity between neurons (Vaccaro *et al*., 2017). Defects in mitochondrial dynamics are a hallmark of many neurological disorders (Sheng and Cai, 2012; Zhu *et al*., 2013; Flannery and Trushina, 2019). To measure mitochondrial motility, we monitored Translocase of Outer Mitochondrial Membrane 20 (TOMM20) fused to a Halo tag for fluorescent labeling. TOMM20 is a member of the TOM complex, a large protein complex spanning the outer mitochondrial membrane (Schleiff *et al*., 1999). We generated iPSCs expressing Halo-tagged TOMM20 using a piggybac transposon and mScarlet as cytoplasmic fill using lentiviral transduction (Methods). To monitor the transport of the mitochondria in axons using live-cell imaging, we used a low-labeling density technique whereby only a fraction of total cells are fluorescently labeled to generate i^3^Neurospheres that are 10% labeled and i^3^Neuron dispersed cultures that are 1% labeled. The lower labeling fraction for the dispersed cultures was needed because of the disorganized meshwork of their axons, in contrast to the organized, radial array in the neurospheres. We validated the selective targeting of exogenous TOMM20-Halo to mitochondria by co-staining for Cytochrome C, a mitochondria localized protein essential for the shuttling of electrons into this organelle (Huttemann *et al*., 2011) (Figure 4B). For i^3^Neurospheres, we identified a field of view that included growth cones (Figure 4A) which, due to the stereotyped radial outgrowth we had previously characterized, ensured the uniform orientation of all axons (Figure 1). In contrast, for live cell imaging of the dispersed i^3^Neurons, we first had to identify the growth cone of each individual neuron to determine axon orientation due to the heterogeneity of axon outgrowth patterns intrinsic to this model. We observed many stationary mitochondria as well as mitochondria exhibiting bidirectional transport, both in the i^3^Neurosphere axons as well as the dispersed i^3^Neuron culture axons (Figure 4C; Supplemental Movie 2). Mitochondrial transport speeds were similar between the dispersed culture model and the neurosphere model at D7 and D11, and consistent with those obtained in an initial characterization of i^3^Neurons (Boecker *et al*., 2020) (Figure 4D). The proportion of motile mitochondria, both in the anterograde and retrograde direction, was higher in the dispersed i^3^Neurons than in the i^3^Neurospheres, ∼ 7.8% *versus* ∼4.6% at D7 and ∼ 4.7% *versus* 4.4% at D11, respectively (Figure 4E). This difference may reflect slight differences in differentiation timelines between the two culture systems. Mitochondrial motility is known to decrease during neuronal maturation (Boecker *et al*., 2020), suggesting that the neurons extending from the neurosphere may be more mature than those in the dispersed culture. The neurosphere assay allowed data acquisition and analysis of 4,308 mitochondria at the D7 timepoint. In contrast, the same acquisition time and data storage space permitted imaging and analysis of only 555 mitochondria in the dispersed culture for the corresponding timepoint, without accounting for the considerable time invested in first identifying axons in the dispersed culture model. Taken together, our data show that mitochondrial motility parameters in i^3^Neurosphere axons are similar to those in dispersed culture and demonstrate the power of the neurosphere model for faster and high-throughput data acquisition for cargo transport analysis in i^3^Neurons.

**Figure 4.**
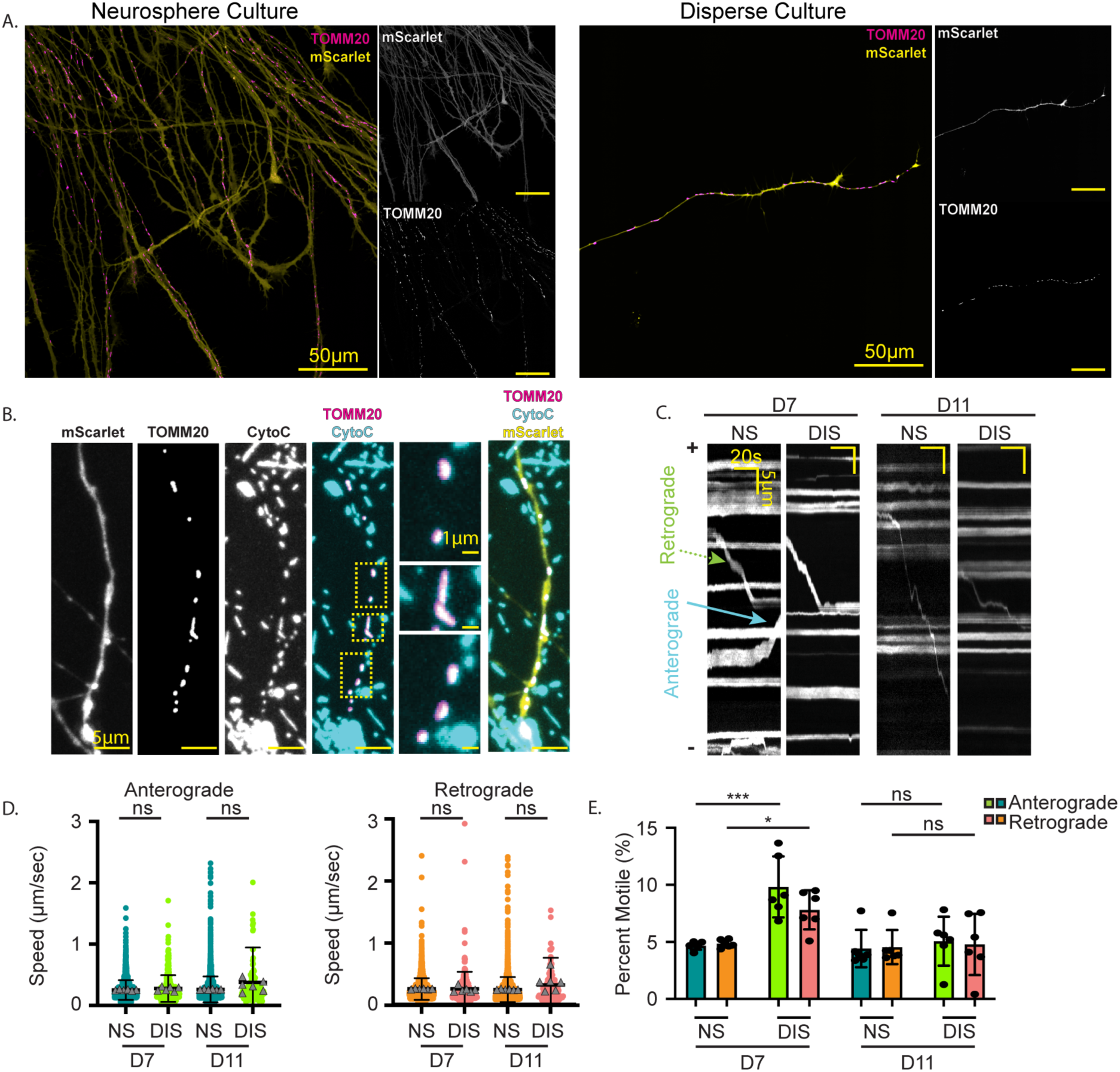
Mitochondrial motility in i^3^Neurosphere axons. (A) i^3^Neurons expressing mScarlet (yellow), TOMM20-Halo labeled with JF646 (magenta), displaying full field-of-view for neurosphere and dispersed cultures. (B) i^3^Neurons expressing mScarlet (yellow), TOMM20-Halo labeled with JF646 (magenta) and stained for cytochrome C (cyan). (C) Kymographs showing mitochondrial motility in i^3^Neurosphere (NS) versus dispersed (DIS) culture at D7 and D11. (D) Anterograde (left) and retrograde (right) average track speed of mitochondria in i^3^Neurosphere axons compared to dispersed i^3^Neuron axons at D7 and D11. Circles represent individual mitochondria, triangles represent mean speed for each individual neurosphere, bars represent mean and S.D.; ns, not significant, *, p < 0.05, **, p < 0.01 by unpaired t-test. D7 NS, N = 4308, D7 DIS, N = 555, D11 NS, N = 5757, D11 DIS, N = 227 tracked mitochondria from six biological replicates from two independent differentiations. (E) Percentage of retrograde and anterograde moving mitochondria. Bars represent mean and S.D., ns, not significant; *, p < 0.05, ***, p < 0.001 by unpaired t-test.

### Longitudinal analysis of synaptic vesicle motility in i^3^Neurosphere axons

Neuronal signaling is driven by synaptic vesicle release from the axon of a presynaptic neuron, and subsequent receptor binding on a postsynaptic neuron. Synaptic vesicles rely on microtubule-based transport from the soma to each synapse, largely mediated by the kinesin-3 motor KIF1A (Okada *et al*., 1995). Synaptophysin (SYP) is a synaptic vesicle transmembrane protein that facilitates membrane fusion for synaptic vesicle release (Kwon and Chapman, 2011). To measure synaptic vesicle motility, we generated iPSCs expressing Halo-tagged SYP using a piggybac transposon and mApple as cytoplasmic fill and delivered it through lentiviral transduction. We then generated i^3^Neurospheres and i^3^Neuron dispersed cultures with low-labeling density (Methods). We validated the targeting of Halo-SYP to synaptic vesicles by immunocytochemistry for synaptophysin (Figure 5A). To facilitate the tracking of motile vesicles against a high background, we photobleached 30 μm regions in the distal axons prior to image acquisition. Without this step, the signal is dominated by brighter, static vesicles, as documented in earlier studies (Dou *et al*., 2024). As expected, labeled synaptic vesicles exhibited both anterograde and retrograde movement in the axons emanating from the core of the i^3^Neurospheres and the dispersed i^3^Neurons (Figure 5B; Supplemental Movie 3). Vesicle transport speeds were consistent across the neurosphere model and the dispersed culture model throughout our time course (Figure 5C).

**Figure 5.**
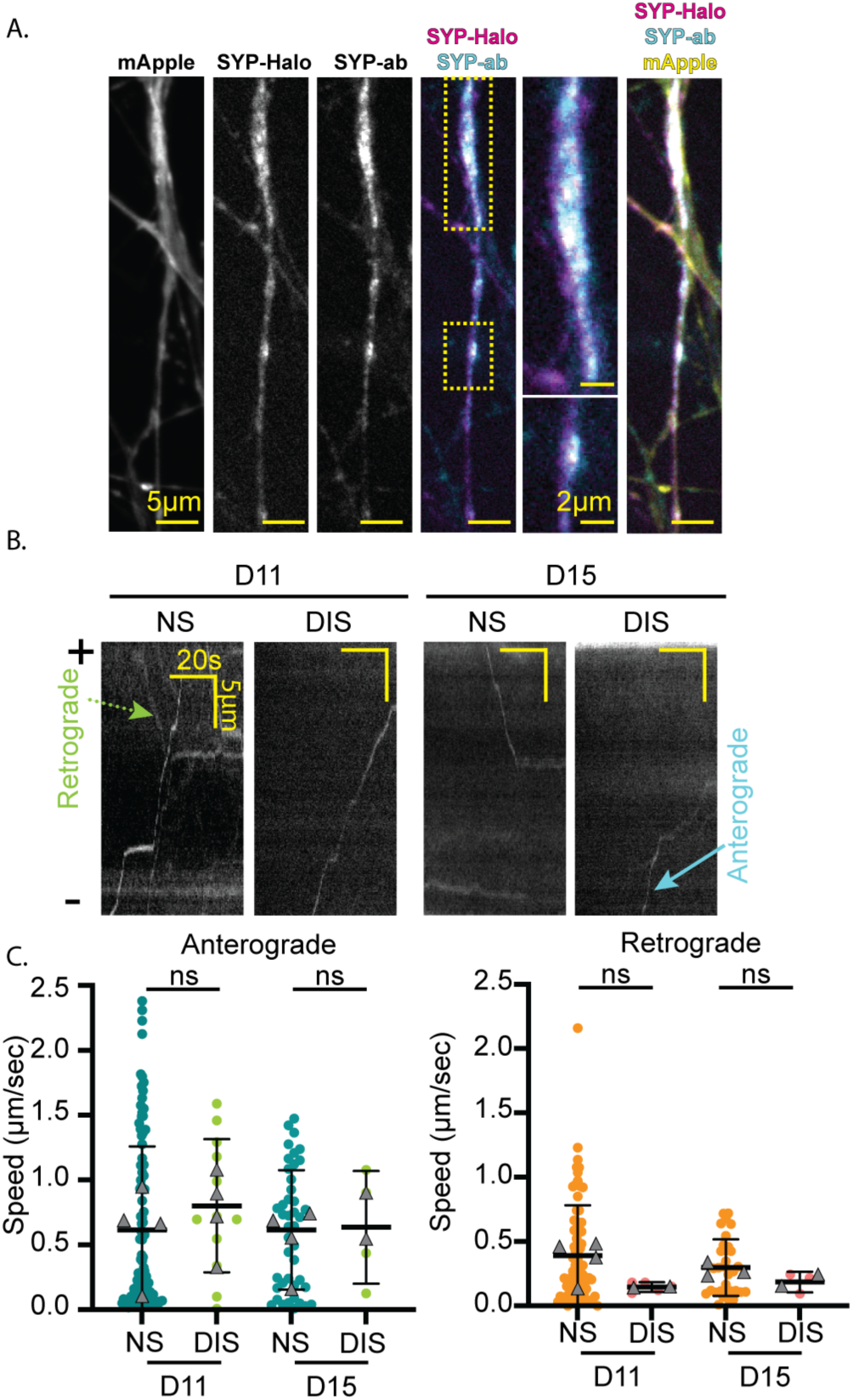
Synaptic vesicle precursor motility in i^3^Neurosphere axons. (A) i^3^Neurons expressing mApple (yellow) and Halo-SYP labeled with JF646 (magenta) and stained for SYP (cyan). (B) Kymographs showing synaptic vesicle motility at D11 and D15. (C) Average speed for anterograde (left panel) and retrograde (right panel) synaptic vesicles in i^3^Neurosphere (NS) axons compared to dispersed (DIS) i^3^Neuron culture axons at D11 and D15. Circles represent individual tracked vesicles, triangles represent mean vesicle speed for each individual neurosphere, bars represent mean and S.D.; ns, not significant; by unpaired t-test. D11 NS, N = 185, D11 DIS, N = 17, D15 NS, N = 75, D15 DIS, N = 7 from four biological replicates from two independent differentiations.

### Longitudinal analysis of lysosome motility in i^3^Neurosphere axons

Lysosomes are essential for proteostasis. They are trafficked throughout axons by kinesin-1 and kinesin-3 motors to maintain homeostasis. Defects in lysosomal motility are a hallmark of neurodegenerative disease (Roney *et al*., 2021). Additionally, lysosomes also assist in transporting other proteins in the axon through cargo adaptors such as the BORC complex (De Pace *et al*., 2024; Ryan *et al*., 2024). To measure lysosomal motility, we used a heterozygous knock-in LAMP1-mEGFP iPSC line. LAMP1 is a trans-membrane protein abundant on the lysosomal membrane (Chen *et al*., 1985). We stably expressed mScarlet as a cytoplasmic fill using lentiviral transduction and then generated neurospheres as well as dispersed cultures with low-labeling density. We validated the selective labeling of lysosomes in the endogenously tagged reporter line by immunocytochemistry for LAMP1. This showed clear co-localization between the antibody and the GFP signal (Figure 6A). As expected, we observed robust bidirectional motility of lysosomes in the neurosphere and dispersed culture axons (Figure 6B; Supplemental Movie 4). Lysosome speed was similar at D7 and D11 for both anterograde and retrograde motility (Figure 6C). Furthermore, at D7 and D11, anterograde and retrograde motility parameters are similar in neurospheres and dispersed cultures.

**Figure 6.**
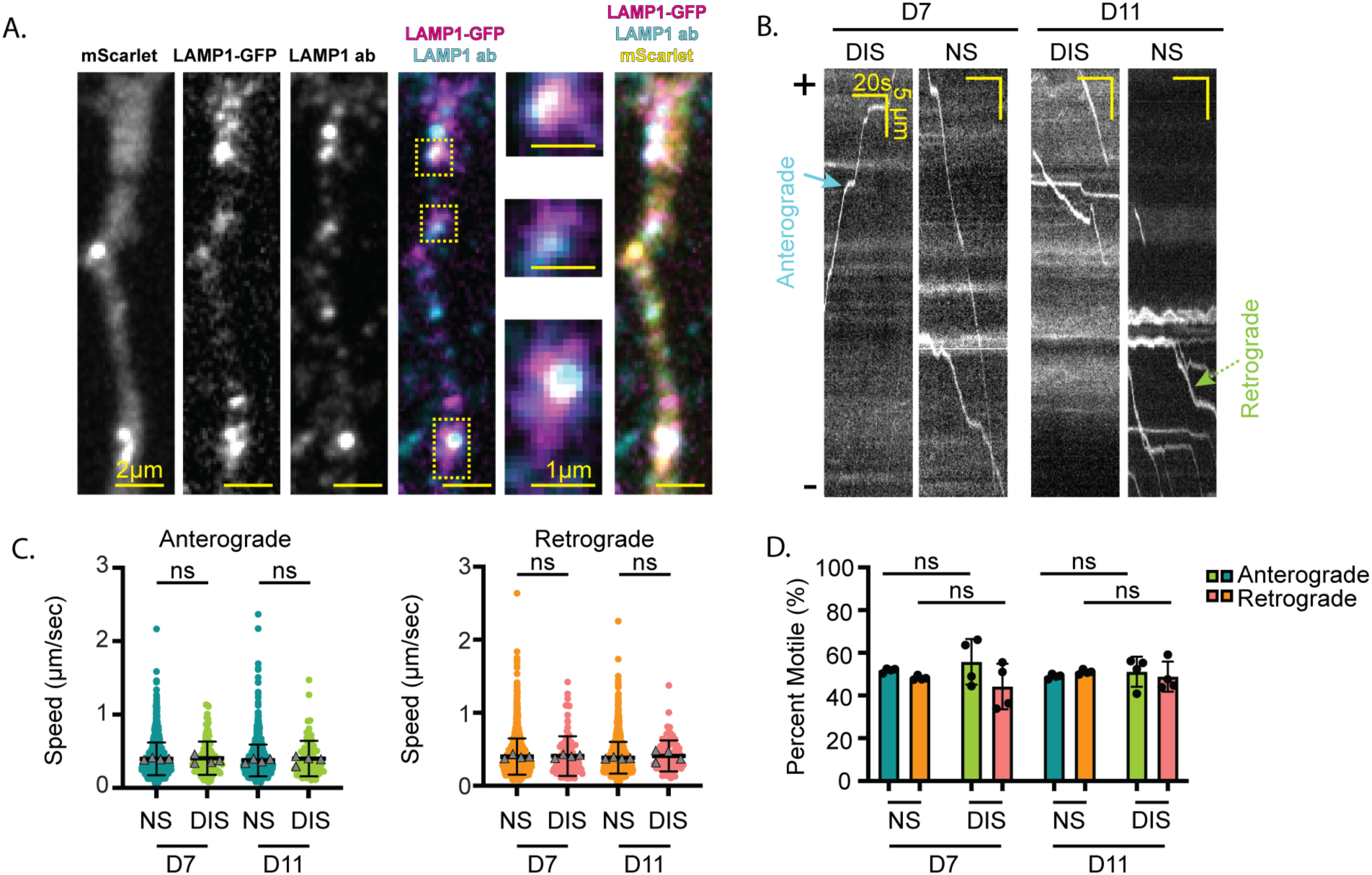
Lysosome motility in i^3^Neurosphere axons. (A) i^3^Neurons expressing mScarlet (yellow), LAMP1-mEGFP (magenta) and stained for LAMP1 (cyan). (B) Kymographs showing lysosomal motility at D7 and D11. (C) Anterograde (left) and retrograde (right) average speed for lysosomes in i^3^Neurosphere (NS) axons compared to dispersed i^3^Neuron (DIS) axons at D11 and D15. Circles represent individual lysosomes, triangles represent mean speed for each individual neurosphere, bars represent mean and S.D.; ns, not significant by Kruskal-Wallis test. D7 NS, N = 3076, D7 DIS, N = 255, D11 NS, N = 2027, D11 DIS, N = 164 lysosomes from four biological replicates from two independent differentiations. (D) Percentage of retrograde and anterograde moving lysosomes. Bars represent mean and S.D., ns, not significant by unpaired t-test.

### An i^3^Neurosphere-glia co-culture model

*In vivo*, neurons interact with glia, which provide a supportive growth environment (Demmings *et al*., 2025), provide metabolites, and regulate neurotransmitters (Ichida and Kiskinis, 2015). Disruptions in neuron-glia interactions contribute to the progression of neurodegenerative diseases such as Alzheimer’s disease (Luchena *et al*., 2022). We therefore established a neurosphere-glia co-culture model that combines all the advantages of the i^3^Neurospheres to investigate i^3^Neuron-glial interactions. To do this, we cultured mouse astrocytes in the imaging dish for two days and then replated the mScarlet- and TOMM20-Halo-labeled i^3^Neurospheres on top of the astrocyte monolayer (Methods; Figure 7A). We first compared axon outgrowth and found that co-cultured i^3^Neurospheres grow in a similar pattern to the monoculture ones, with axons growing radially outwards from the center. Consistent with the trophic support provided by the glial substratum, axonal outgrowth is slightly faster and reaches longer distances (Figure 7B). We note that at later timepoints co-cultured neurospheres grow with less uniformity than the monoculture neurospheres (Figures 7B, C and 1C). This could be due to developmentally timed interactions with the glial substratum or adherence of neurites to the glial monolayer. We then imaged mitochondrial motility in the i^3^Neurosphere-glia co-culture. This showed that anterograde and retrograde motility of mitochondria in the co-cultured i^3^Neurospheres are similar to those in isolated monocultured i^3^Neurospheres at D7, D11, and up to D20 (Figure 7D). Additionally, analysis shows a similar proportion of mitochondria moving in the anterograde and retrograde direction as in the monocultures (Figure 7E and 4E). Our data show that i^3^Neurospheres can be adapted to a glia co-culture model, facilitating the exploration of questions related to the impact of neuron-glia interactions on axonal outgrowth, cytoskeletal regulation and organelle dynamics.

**Figure 7.**
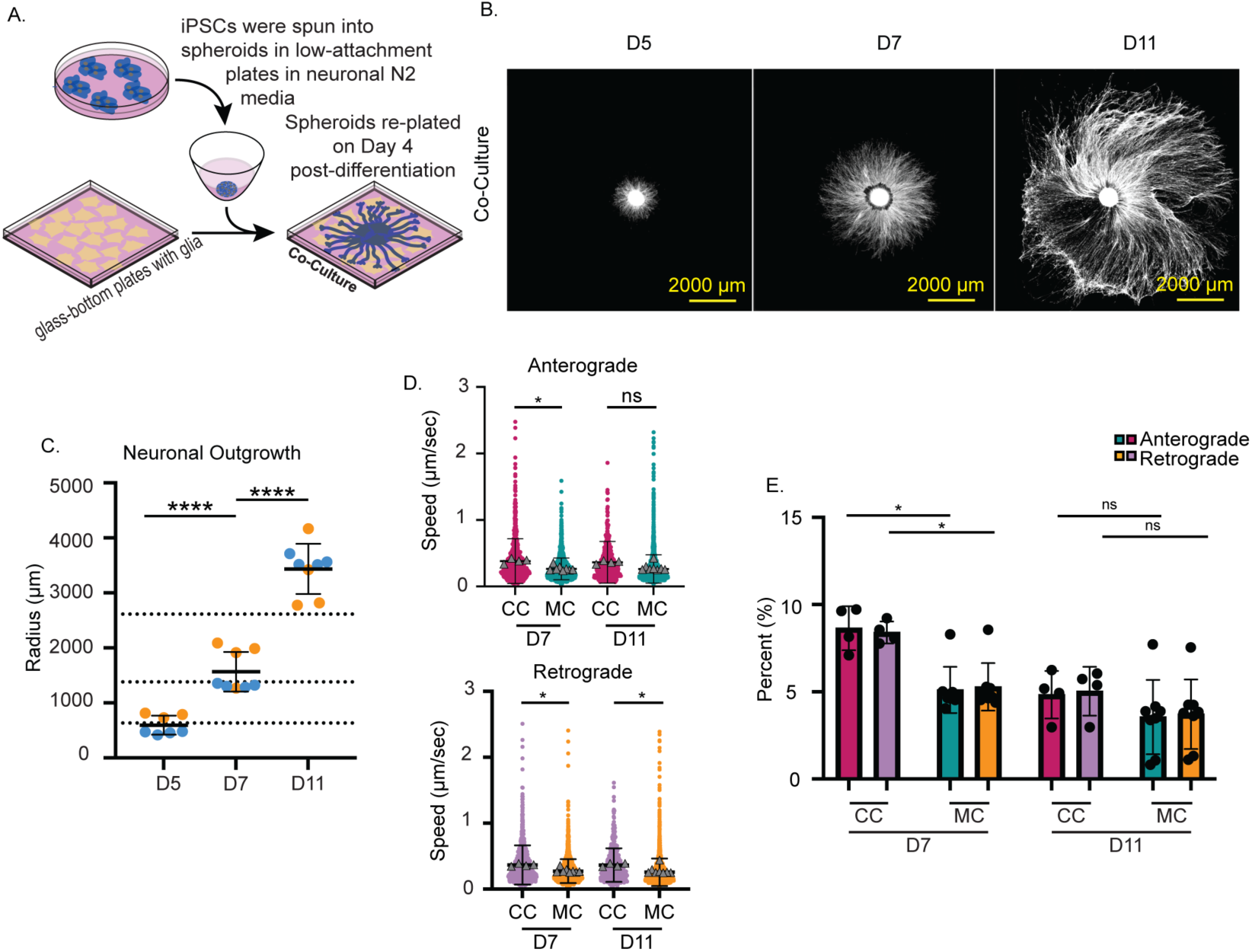
I^3^Neurosphere co-culture with glia. (A) Schematic showing i*^3^*Neurosphere generation and replating onto a bed of mixed astrocytes. (B) i^3^Neurosphere outgrowth at D5, D7, and D11. (C) Average neurosphere radius at D5, D7, and D11. Circles represent individual neurospheres, different colors represent independent differentiations. Bars represent mean and S.D., **, p<0.01 by unpaired t-test. N= 7, 8, and 8 for D5, D7 and D11 neurospheres, respectively from two independent differentiations. Dotted lines indicate the average outgrowth from the monoculture neurospheres at the equivalent time point. (D) Anterograde (top) and retrograde (bottom) average speed of mitochondria in i^3^Neurosphere axons cultured as monoculture (MC) compared to those co-cultured with mouse glia (CC) at D7 and D11. Circles represent individual tracked mitochondria, triangles represent mean speed for each individual neurosphere. Bars represent mean and S.D., ns, not significant; ****, p < 0.0001 by Kruskal-Wallis test. D7 CC, N = 11501, D7 MC, N = 8517, D11 CC, N = 10082, D11 MC, N = 11249 tracked mitochondria from two independent differentiations for CC, and three independent differentiations for MC. (E) Percent of motile mitochondria defined as the proportion of total particles moving more than 1μm of total distance. Bars represent mean and SD.; ns, not significant, *, p < 0.01 by unpaired t-test. MC, monoculture, CC, co-culture with mouse glia.

## Discussion

Human iPSC-derived neurons are a powerful tool to study cytoskeletal dynamics and cargo trafficking, and the dysregulation of these processes in disease. The visualization of these processes in individual axons using dispersed neuron cultures is challenging because of the unique neuronal morphology and asynchrony of axonal outgrowth. Axonal processes are long and often overlap in dispersed culture, making identification, isolation and analysis of these processes nontrivial. Existing solutions involve the use of extremely sparse labeling to allow isolation of individual processes; however, this method limits data set size by increasing the time needed to identify axons, as well as image acquisition time and data storage since a single image or movie needs to be collected for each individual neuron. More efficient, compartment-specific imaging requires microfluidic devices; however, these limit imaging to only the axonal sub-compartment, inhibit axonal specific imaging early in differentiation and limit the data acquired to the number of microfluidic channels available. Microfluidic devices are also difficult and time-consuming to maintain and can be expensive to manufacture. The primary neuronal explant model is an outstanding method for studying thousands of axons that project radially from a spheroid body; however, it does not allow study of human neurons and therefore is subject to all the technical and model-related limitations therein. While spheroid cultures of human neurons have been put forth as potential platforms for high-throughput developmental neurotoxicity screening, these existing “neurosphere assays” rely on time-consuming generation from embryoid bodies, hNPCs or hiPSCs using small molecule-mediated differentiation protocols that ultimately yield a heterogenous mix of astrocytes, oligodendrocytes, and neurons (Moors *et al*., 2009; Bamba *et al*., 2016; Schmuck *et al*., 2017; Koch *et al*., 2022). Despite the observation of a general radial array of neurites, this model has never been adopted for stereotyped spatial designation of neuronal compartments or for higher-resolution live-cell imaging applications because of cell-type heterogeneity, including confounding radial glia fibers which promote dispersion of neurons out of the central sphere. Here, we characterize and validate a neurosphere model comprised of a homotypic population of i^3^Neurons as a cost-effective and technically feasible model for the study of microtubule function in human iPSC-derived i^3^Neurons and their axons. Our i^3^Neurospheres model system combines high-resolution high-throughput imaging with facile live-cell reporter line generation and genetic engineering in i^3^Neurons to provide the research community a powerful tool to study cytoskeletal dynamics and other processes in the axons of human neurons.

We show the feasibility of rapid and easy data collection on process outgrowth for thousands of i^3^Neurons in neurospheres due to their stereotyped radial outgrowth pattern. The large datasets obtained ensure rigor and reproducibility. They also allow the detection of small effects that can be physiologically relevant, especially for understanding age-related diseases and the more subtle effects of disease mutations. We show that the outer section of the i^3^Neurosphere is comprised of axons: five days beyond neuronal induction, all growth cones and neurites within the field of view of the distal perimeter of the neurosphere are MAP2-negative, and positive for axon-specific neurofilaments, confirming axonal identity. Furthermore, as expected for axons, these processes exhibit increasingly unipolar, plus-end-out microtubule growth, further confirming axonal differentiation. The stereotyped radial pattern of axonal outgrowth in the i^3^Neurospheres allows easy identification of growth cones and facilitates parallel imaging in many axons as a function of distance from the growth cone. This assay configuration can therefore support the identification of regulators of axonal outgrowth and any investigation into axon pathfinding, and how disease-related proteins alter axon guidance.

Microtubule-based cargo transport is one of the most fundamental and studied neuronal processes. I^3^Neurons were previously shown to have microtubule-dependent cargo transport parameters similar to those recorded in primary neurons (Boecker *et al*., 2020). Our analysis, in turn, shows that i^3^Neurosphere cargo transport parameters exhibit great similarity to the low-density dispersed i^3^Neurons previously benchmarked, while providing significant advantages for rapid data collection. I^3^Neurosphere and dispersed culture axons showed no significant differences in the speed of mitochondrial, lysosomal, and synaptic vesicle transport. The directionality of transport for mitochondria, lysosomes, and synaptic vesicles also showed minimal variation between the two model systems. We also show that the neurosphere model can be easily adapted to a co-culture system with glia. Glia co-culture allows neurons to be cultured for longer times and in a more physiological context, important for disease modeling, while also making possible the study of non-cell autonomous biology. Depending on the nature of the question, i^3^Neurospheres can be cultured for longer times than the ones described in our study. This can be achieved by using imaging-compatible culture dishes with larger-diameter wells (our study uses 1 cm^2^ diameter wells) to prevent axons from growing beyond the boundaries of the wells, which would result in the loss of the radial orientation critical for high-throughput, high-resolution imaging. The use of these culture dishes will increase the cost of experiments but will not introduce any experimental difficulties.

One current limitation to the neurosphere model is that the axons in their radial configuration are spatially constrained from forming synapses. This could be resolved by culturing neurospheres on top of unlabeled dispersed cultured neurons, which would provide underlying dendrites for synaptic connectivity. Alternatively, Holzbaur and colleagues recently developed an innovative way for i^3^Neurons to form robust pre-synapses using exogenous expression of neuroligin-1 (NL1) in HEK293 cells (Aiken and Holzbaur, 2024). Co-culturing neurospheres with NL1-expressing HEK293 cells, or on micropatterned HEK293 cells would also enable engineered synapse formation *in vitro* while maintaining the advantages the i^3^Neurospheres provide.

In conclusion, we present an advantageous model system for studying human axons at scale, and provide in-depth characterization of axonal outgrowth, neurite identity, microtubule dynamics, and microtubule-based transport parameters in this format. We anticipate that our work will be a useful reference study and toolbox for the field to study microtubule-dependent and other neuronal processes in neurobiology and human disease.

## Author Contributions

S.A.C.W. generated cell lines, performed experiments, analyzed data and made figures; E.D.M. and S.L.S generated genetic constructs, generated cell lines, provided and supervised experimental methodology; M.S.F. devised neurosphere methodology; M.E.W. contributed cell lines, genetic constructs; E.D.M., S.L.S. and A.R.M. devised project, supervised work and aided in interpretation of results; S.A.C.W. and E.D.M. wrote first draft; S.A.C.W., E.D.M., S.L.S. and A.R.M. edited manuscript; all authors read and approved manuscript.

## Acknowledgements

We thank J. Spector (National Institute of Neurological Disorders and Stroke, NINDS) for imaging help and creating imaging analysis pipelines, S. Anderson, M. Kirby from the Flow Cytometry core (National Human Genome Research Institute) for cell sorting. This work was partially supported by a Director’s Innovation Challenge Award to A.R.M., a fellowship from National Institute of Neurological Disorders and Stroke (NINDS) to E.D.M., a P.R.A.T; fellowship to S.L.S. from the National Institute of General Medical Sciences. A.R.M. is supported by the intramural program of NINDS. The contributions of the NIH authors were made as part of their official duties as NIH federal employees, are in compliance with agency policy requirements, and are considered Works of the United States Government. However, the findings and conclusions presented in this paper are those of the authors and do not necessarily reflect the views of the NIH or the U.S. Department of Health and Human Services.

## Methods

### Human i^3^Neuron Dispersed Culture and Differentiation

The iPSC line termed “i11wmNC” was used for experiments. This line was previously derived from the well-characterized WTC11 line (Miyaoka *et al*., 2014), contains a dox-inducible NGN2 stably integrated at the AAVS1 safe-harbor locus, and contains dCas9-BFP-KRAB stably integrated at the CLYBL safe-harbor locus (Wang *et al*., 2017; Fernandopulle *et al*., 2018). Only experiments measuring lysosomal motility used a LAMP1-mEGFP line that was not derived from i11wmNC, but it was derived from the WTC11 background. See “Cell Line Generation” below.

Cells were cultured and differentiated as described previously (Fernandopulle *et al*., 2018). iPSCs were cultured on plates coated with Matrigel (VWR #BD354277). Culture media StemFlex (Gibco #A3349401) was changed every 1-2 days. To differentiate iPSCs into neurons, iPSCs were washed with Phosphate-buffered saline (PBS) to remove debris and incubated with 1mL Accutase (Gibco #A1110501) at 37°C in 5% CO_2_ for 5 minutes. Cells were collected in a solution of 1mL StemFlex and 2mL PBS and centrifuged at 300rcf. After centrifugation, cells were resuspended in 1mL neuronal induction media N2 (containing KO-DMEM/F12 (Gibco #12660012), 1% N2 supplement (Gibco #17502048 100x), 1% NEAA (Gibco #11140050), 1% GlutaMAX (Gibco #35050061), 50nM Chroman-1 (MCE #HY-15392), and 0.002mg/mL Doxycycline (Sigma #D9891)) and plated at a density of 1 million/well on a Matrigel-coated 6-well plate. On Day 3, cells were incubated with 1mL Accutase at 37°C for 5 minutes. After collection and pelleting, supernatant was removed, and cells were resuspended in BrainPhys Neuronal Media (StemCell Tech, #05790) supplemented with 2% B27+ supplement (Gibco #A35828_50X), 0.01 μg/mL BDNF (Peprotech #450-02), 0.01μg/mL NT-3 (Peprotech 450-03), 0.0012mg/mL Laminin (R&D Systems #3446-005-01_6mg/ml), 0.002mg/mL Doxycycline (Sigma #D9891), notated as BrainPhys+. To prepare for replating, glass-bottom 8-well ibidi imaging dishes were coated with 0.1mg Poly-L-Ornithine (PLO) in water with 10% borate buffer (100mM boric acid, 75mM NaCl, 25mM sodium tetraborate, pH 8.4) by overnight incubation at 37°C. PLO coated wells were washed 4x with water, and air dried. After drying, plates were incubated with 15μg/mL laminin (R&D Systems #3446-005-01) in PBS at 37°C for 2 hours. After coating, laminin solution was removed and 250μL BrainPhys+ was added to each well. Cells underwent gentle half-media changes of BrainPhys+ Neuronal Media every 2-3 days.

### I^3^Neurosphere Generation

I^3^Neurospheres were generated from the same cell lines as the dispersed cultures for each experiment. iPSCs were dissociated with Accutase, pelleted, and resuspended in the same manner as above. For each neurosphere, 10,000 cells were suspended in 30μL of N2 neuronal induction media and plated in low-attachment 384-well round-bottom plates (Corning, 12456722) after pre-washing wells once with N2 media. Plates were then spun at 500 rcf (Eppendorf 5810, A-4-62 Rotor with MWP buckets) for 10 minutes and inspected to ensure cell aggregation in the center of the well. The plate was then incubated at 37°C in 5% CO_2_. 24 hours later (Day 1 post-differentiation), 80μL of N2 media was added to the wells. On Day 4, 40μL of media was removed from each well, neurospheres were picked up using a wide orifice 200μL pipette tip (Rainin #30389241) and carefully placed into the center of a well coated with PLO and laminin (as described above) containing 250μL of BrainPhys+ neuronal media for monoculture. For co-culture, astrocyte media was aspirated from the well and 250μL of BrainPhys Neuronal Media was added. Neurospheres were lifted from the 384-well plate using a wide-orifice 200μL pipette tip and placed gently in the center of each well. All neurospheres (mono-culture and co-culture) underwent extremely gentle half-media changes every 2-3 days using BrainPhys+ neuronal media.

### Cell Line Generation

For neurite outgrowth assays, i11wmNC cells expressing cytoplasmic mScarlet and EB1-EGFP were used. For microtubule dynamics assays, cells expressed cytoplasmic mScarlet and EB1-EGFP. To obtain a clonal EB1-EGFP line, cells were plated at a very low density (1,000 cells in a 10 cm dish) and grown for 7-10 days before manually picking isolated clones using Lumascope (EtaLuma). A single, homogenous, population of cells was picked and cultured based upon expression of both green and red fluorescence.

To generate the TOMM20-Halo and the Halo-SYP stable lines, the following piggybac transfection protocol was performed. 800,000 iPSCs were plated into one well of a matrigel-coated 6 well plate in Essential 8 medium (Life Technologies #A1517001) supplemented with 50nM chroman-1 (MedChem Express HY-15392), 5mM Emricasan (Selleckchem #S7775), Polyamine supplement (Sigma #P8483) diluted 1:1000 for a final amine concentration 0.423 μg/mL, 0.7mM Trans-ISRIB (Tocris #5284), termed “CEPT”. Transfection mixture was prepared in an Eppendorf tube using OptiMem (Life Technologies #31985070), 1 μg of Halo-SYP or TOMM20-Halo DNA, 1 μg of piggybac transposase and 6mL of LipoStem (Life Technologies #STEM00003). The transfection mixture was mixed thoroughly and incubated at room temperature for 30 minutes. Immediately before adding transfection mixture, fresh Essential 8 supplemented with CEPT was applied to cells. Transfection mixture was added to cells 1 hour after plating. The next day transfected cells were replated into a 10 cm plate. Cells were then FACS-purified for positives, specifically low-expressing positives to avoid artifacts of hyper-overexpression. TOMM20-Halo was introduced into the mScarlet/EB1-EGFP i11wmNC iPSC line. Halo-SYP was introduced into the i11wmNC iPSC line. Later, Halo-SYP cells were infected with lentiviral cytoplasmic mApple and FACS purified for co-expression. LAMP1-mEGFP cells are a heterozygous knock-in iPSC line obtained from the Allen Institute and generated in the WTC11 genetic background (Cell line ID: AICS-0022 cl.37). LAMP1-mEGFP cells were transfected with 1 μg of a piggybac-NGN2 plasmid encoding hygromycin resistance according to the same piggybac protocol described above. These cells were additionally transduced with a cytoplasmic mScarlet.

### Immunocytochemistry (ICC)

Culture media was removed from wells, and 200μL of PBS supplemented with 4% Paraformaldehyde (PFA) was added to each well. Samples were incubated at 37°C for 15 minutes. After incubation, samples were washed 3 times with 200μL PBS. Samples were then permeabilized in 200μL PBS with 0.2% Triton for 10 minutes. Samples were then blocked in 200μL PBS with 2% Bovine Serum Albumin (BSA) for 15 minutes. Primary antibodies were diluted in 1% BSA. 200μL primary antibody solution was added to each well and incubated at 4°C overnight. Primary antibody was removed the next day, and samples were washed carefully three times with PBS. Fluorophore-conjugated secondary antibodies were diluted in 1% BSA, added to each well, and incubated at 37°C for 1 hour. Secondary antibody was removed and samples were washed three times. Cells were stored at 4°C in PBS for imaging. Primary antibody dilutions were as follows: ch-MAP2 (Abcam #ab5392, 1:5000), m-Tau (Millipore #MAB3420, 1:1000), pan-axonal SMI-312 (BioLegend 837904, 1:500), m-Cytochrome C (BD Biosciences #556432, 1:500), m-SYP (Millipore Sigma #S5768, 1:500), and rb-LAMP1 (Proteintech #21997-1-AP, 1:500). Fluorophore-conjugated secondary antibodies used were anti-m Alexa 488 (Invitrogen #A21202), anti-ch Alexa 646 (Invitrogen #A78948), anti-m Alexa 405 (Invitrogen #A48257), and anti-rb Alexa 405 (Invitrogen #A48258), all diluted 1:500.

For MAP2/tau staining, i^3^Neurospheres were fixed on D5, D7, D9, and D11 post-differentiation. For TOMM20-Halo, Halo-SYP, and LAMP1-mEGFP ICC validation, cells were fixed at D20. For TOMM20-Halo and Halo-SYP, samples were incubated for 15 minutes prior to imaging with 10nM Janelia Fluor 646 HaloTag Ligand (JF646) (Promega). For pan-axonal neurofilament staining, i^3^Neurospheres were fixed at D5 and permeabilization was changed to 0.1% Triton for 20 minutes.

### Glia Culture

Mouse C57 Brain Mixed Astrocytes (Lonza #MASM330) were cultured on plates coated with Matrigel in culture media of AGM Astrocyte Growth Medium (Lonza #CC3186). For co-culture with iPSC-derived neurons, mixed astrocytes were plated at a density of 42,000 cells in the center of one well of an 8-well glass-bottom ibidi coated with matrigel and air-dried. Media was changed every 2 days while preparing for neuronal cells.

### Live Imaging

Live imaging was performed on a Nikon Ti2 with a CSU-W1 spinning disk confocal head using a Hamamatsu Fusion BT camera and Nikon LU-NF laser system with 405nm, 488nm, 561nm, and 647nm solid-state diode lasers. All live imaging used an OKO labs live imaging chamber at 37 °C and 5% CO_2_.

Neuronal outgrowth imaging was performed live and tiling was performed as needed to capture all projections. On days 5 and 7 post-differentiation, a 10x/0.3NA objective (Nikon, Plan Fluor 10x OFN25 DIC N1) was used and z-stacks were 12 μm in height with a 3μm step size. On day 11, a 4x objective (Nikon, Plan 4x) was used and z-stacks were 30μm in height with a 5μm step size.

Live imaging EB1-EGFP comets in distal axon regions (200 μm or less from the growth cone) was performed using a 60x-H_2_O immersion objective (Nikon, Plan Fluor 40x Ph2 DLL) Time lapse images were collected using the 488nm laser line with a 2 second framerate for a duration of 60 seconds.

For live imaging of TOMM20-Halo and Halo-SYP, samples were incubated for 15 minutes prior to imaging with 10nM JF646. TOMM20-Halo live imaging was performed using a 60x-H_2_O immersion objective in distal axon regions (within 200μm of the growth cone). One still image of mScarlet expression was captured using the 561nm laser line followed by time lapse images of JF646-labeled TOMM20-Halo using the 647nm laser line with a 0.5 second framerate for 60 seconds.

Halo-SYP live imaging was performed using a 60x-H_2_O immersion objective. Distal axon regions (within 200μm of the growth cone) were photobleached using a 405nm CyrLas solid-state laser with 30μs of dwell time for 60 seconds. After photobleaching, one still image of mApple expression was captured using the 561nm laser line followed by a time lapse images of JF646-labeled Halo-SYP captured using the 647nm laser line with a 0.5 second framerate for a duration of 60 seconds.

LAMP1-mEGFP live imaging was performed using a 60x-H_2_O immersion objective in distal axon regions (within 200μm of the growth cone). One still image of mScarlet expression was captured using the 561nm laser line followed by time lapse images of LAMP1-mEGFP using the 488nm laser line with a 0.5 second framerate for a duration of 60 seconds.

### Imaging of fixed samples

I^3^Neurospheres were imaged using a Nikon Ti2 with a CSU-W1 spinning disk confocal head using a Hamamatsu Fusion BT camera and Nikon LU-NF laser system with 405nm, 488nm, 561nm, and 647nm solid-state diode lasers. For Tau/MAP2 imaging, a 10x objective (Nikon, Plan Fluor 10x OFN25 DIC N1) was used and tiling was performed as needed to capture all projections. Z-stacks were 60μm in height with a 6μm step size. For cargo labeling validation, a 60x-H_2_O objective was used. Z-stacks were 7μm in height with a 1μm step size. For TOMM20-halo and Halo-SYP validation, samples were incubated for 15 minutes with 10nM JF646 prior to fixation as mentioned previously.

### Image Analysis and Quantification

For neuronal outgrowth analysis, images of neurospheres were tiled, max-projected, and run through an analysis pipeline in NIS Elements. Briefly, the analysis used thresholding to segment the inner sphere (the somatodendritic core) and the outer sphere (the periphery of neurite outgrowth) based on signal intensity (Figure 1D). Segmented shapes were analyzed for area, the radius was calculated from the “equivalent diameter” of a circle with the same area. Inner sphere radius was subtracted from outer sphere radius to control for slight variations in cell numbers in the neurosphere core. For MAP2/tau analysis, intensity plots were generated in Fiji by taking a radial line scan from the center of the sphere outwards, rotated every 1 degree to generate an averaged intensity of the signal relative to distance from the center. For EB1-EGFP comets, the percent anterograde comets was determined by generating kymographs using the MultiKymograph macro in Fiji along 26 μm-long sections of neurite and counting numbers of anterograde and retrograde tracks in each kymograph. For microtubule growth speed analysis, sections of growth cone in each image were cropped and each cropped image was analyzed using TrackMate (Tinevez *et al*., 2017; Ershov *et al*., 2022). For TOMM20 and LAMP1 particle motility, particles were analyzed using the TrackMate plugin in Fiji. An ROI was drawn at the minus-end for a given field of view and the ‘Distance to ROI’ feature was used to determine directional motility. Motile particles were distinguished from stationary particles by a net displacement greater than 1μm, and mean speed and fraction of motile particles was plotted. For SYP motility, kymographs were drawn from a photobleached section of the neurite using NIS Elements. Tracks were traced in Fiji and the coordinates of each track collected. Mean speed was calculated by taking the slope between each frame and averaging.

---

**Figure S1 (related to Figure 2).**
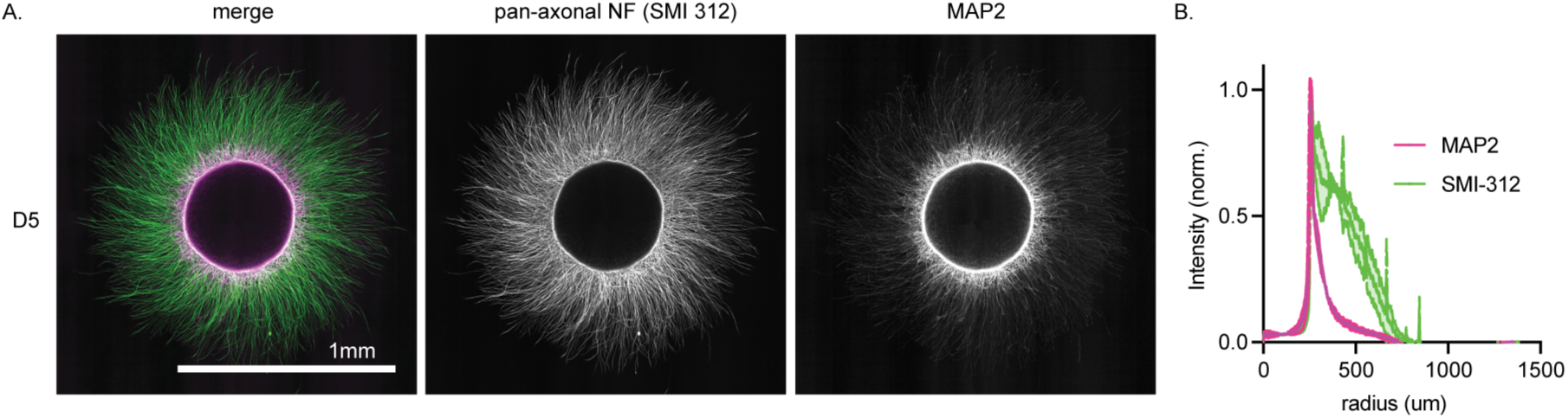
Additional neuronal polarity markers in i^3^Neurospheres define dendritic and axonal compartment boundaries. (A) Representative image of a D5 neurosphere immunolabeled for MAP2 (magenta) and pan-axonal neurofilament marker SMI-312 (green). (B) Line plots along the neurosphere radius (from center to periphery) showing average intensity of MAP2 (magenta) and SMI-312 (green) signal, normalized to their respective highest intensity as in Figure 2B; N = 2 neurospheres.

**Movie S1 (related to Figure 3). Microtubule dynamics in i^3^Neurosphere axons.** Representative video of EB1-GFP microtubule comets in distal i^3^Neurosphere axons. i^3^Neurons are co-expressing mScarlet (visible in initial frames), and red boxes delineate the inset regions magnified in subsequent frames.

**Movie S2 (related to Figure 4). Mitochondrial motility in i^3^Neurosphere axons.** Representative video of mitochondrial motility in distal i^3^Neurosphere axons using JF646-labeled TOMM20-Halo. i^3^Neurons are co-expressing mScarlet (visible in initial frames), and the red box delineates the inset regions magnified in subsequent frames.

**Movie S3 (related to Figure 5). Synaptic vesicle precursor motility in i^3^Neurosphere axons.** Representative video of synaptic vesicle precursor motility in distal i^3^Neurosphere axons using JF646-labeled Halo-SYP. i^3^Neurons are co-expressing mScarlet (visible in the initial frames), the red box delineates the inset regions magnified in subsequent frames, the pink box delineates the photobleached region magnified in subsequent frames.

**Movie S4 (related to Figure 6). Lysosomal motility in i^3^Neurosphere axons.** Representative video of lysosomal motility in distal i^3^Neurosphere axons using a haploid knock-in LAMP1-mEGFP cell line. i^3^Neurons are co-expressing mScarlet (visible in the initial frames), the red box delineates the inset regions magnified in subsequent frames, the pink box delineates the inset of the photobleached region magnified in subsequent frames.

